# Resolving tricky nodes in the tree of life through amino acid recoding

**DOI:** 10.1101/2022.02.24.479670

**Authors:** Mattia Giacomelli, Maria Eleonora Rossi, Jesus Lozano-Fernandez, Roberto Feuda, Davide Pisani

**Affiliations:** School of Biological Sciences, University of Bristol, Life Sciences Building, Tyndall Avenue, Bristol, BS8 1TQ UK; School of Earth Sciences, University of Bristol, Life Sciences Building, Tyndall Avenue, Bristol BS8 1TQ, UK; Department of Genetics, Microbiology and Statistics, Faculty of Biology, University of Barcelona, Spain; Department of Genetics and Genome Biology, University of Leicester, Leicester, UK

## Abstract

Genomic data allowed for a detailed resolution of the tree of life. Yet, tricky nodes such as the root of the animal, plants, eukaryotes, bacterial and archaeal trees remain unresolved. Genomic datasets are heterogeneous as genes and species evolve under different selective pressures, impending the efficacy of evolutionary analyses. Amino acid recodings were developed to reduce heterogeneity, but clear evidence to justify their use is missing. We use simulated genomic-scale datasets and show that recodings can substantially improve phylogenetic accuracy when tackling tricky nodes. We apply our findings to address the root of the animal tree where the debate centers on whether sponges (Porifera) or comb jellies (Ctenophora) branched out first. We show that results from real data follow predictions from simulated data and indicate that a placement of the ctenophores as the first branching animal lineage is most likely artifactual.

## Introduction

Understanding of the tree of life has enormously increased since genome-scale (i.e. phylogenomic) datasets became available (e.g. (*1–4*)). Yet, some tricky nodes remain that are proving difficult to resolve, such as the root of the bacterial, archaeal, eukaryotes, plants and animal trees (*5–11*). These uncertainties hinder understanding of broader evolutionary questions. For example, uncertainty in animal phylogeny, where the debate centers on whether sponges (Porifera) or comb jellies (Ctenophora) are sister to all other animals (*5, 12*– *19*), affects our understanding of the evolution of body plans, with ramifications reaching areas as different as developmental biology and palaeoecology (*20, 21*).

One reason for the difficulty in resolving tricky nodes is the heterogeneity of the evolutionary process. Genome-scale datasets are heterogeneous as different genes, sites and species evolve under different constraints (*22*). Heterogeneity primarily manifests itself in the form of across-site and -lineage rate heterogeneity (i.e. how many amino acid substitutions are observed at different sites or along different lineages), and across-site and -lineage compositional heterogeneity (i.e. which amino acids are observed at different sites and in different taxa). Evolutionary models that can better account for these forms of heterogeneity (*23–25*) fit the data better. Yet, even models that can take into consideration different forms of heterogeneity, like the CAT-GTR+G model, which accounts for across-site compositional heterogeneity (the CATegory component (*23*)), amino acid replacement rate heterogeneity (the General Time Reversible (GTR) matrix (*26*)) and across-site rate heterogeneity using a Gamma (+G (*27*)) distribution, frequently fail to fit real world datasets (*4, 5*).

Amino acid recodings were developed to attempt reducing compositional heterogeneity (*28–31*). These methods (hereafter also referred to as exchangeability-informed recodings, or just recodings), partition the 20 amino acids into bins based on how frequently they exchange with each other (*5*) – details in Fig.1. Exchangeability-informed recodings are generalizations of standard masking protocols, such as the amino acid recoding of nucleotide data (*32*). When exchangeability-informed recodings are used, substitutions among amino acids with similar chemical and physical properties are synonymized (Fig. 1). This is analogous to the process by which codons are synonimized when nucleotides are recoded into amino acids. The rationale underpinning the synonymization procedure is that substitutions among amino acids with similar properties (the within-bin substitutions – Fig. 1) are expected to have a smaller negative impact on the fitness of the coded protein when compared against substitutions among amino acids with different properties (across-bin substitutions – Fig.1, (*5*)). Within-bin substitutions are thus expected to be more abundant and less constrained, contributing more heterogeneity to the data.

**Figure 1.**
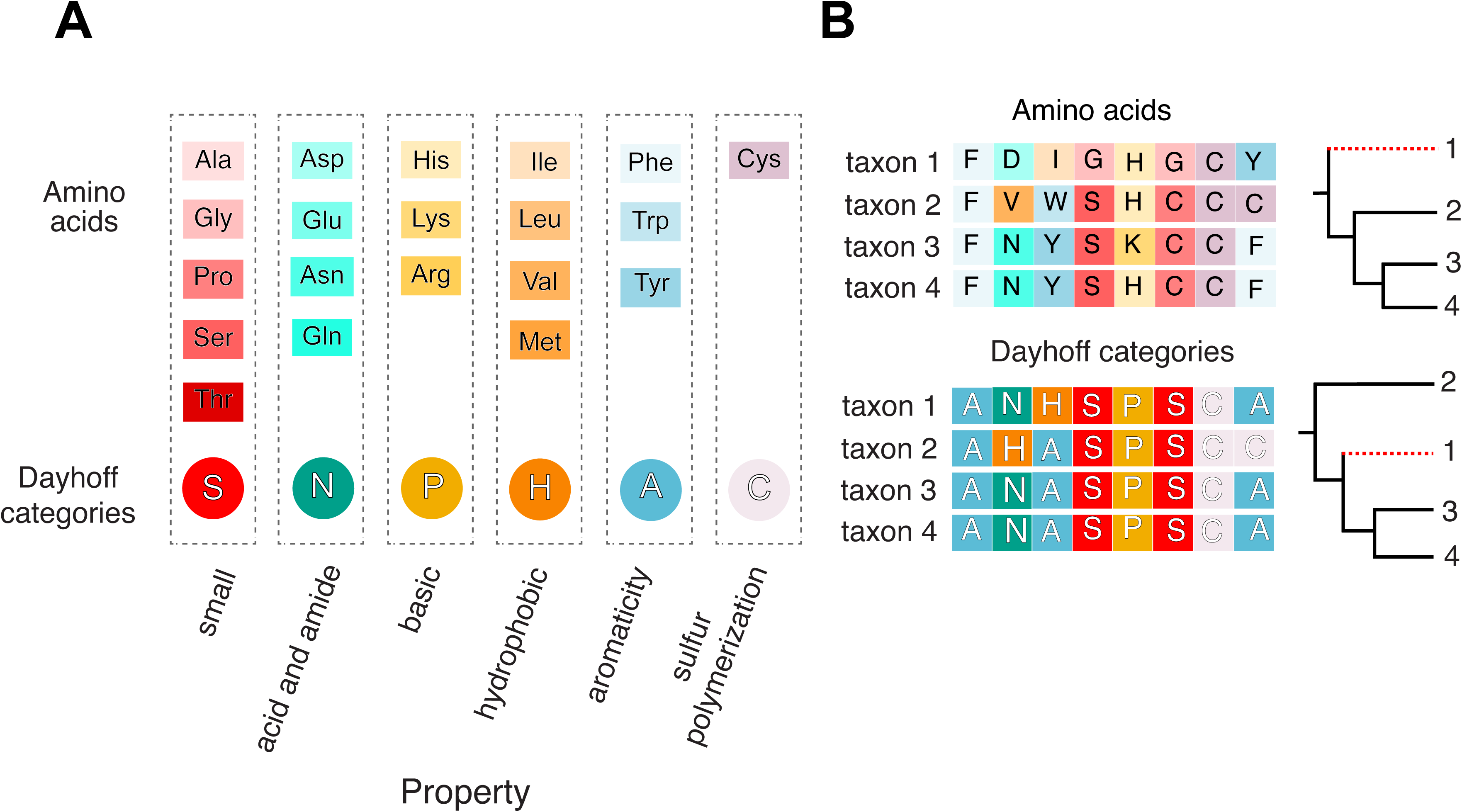
The six-bin Dayhoff-6 recoding (*60*) is the most widely used exchangeability- informed, and is the one primarily tested in this study. Other recodings include the 6-bin KGB6 (*30*), and SR6 (*31*), see (*5*). These recodings differ from Dayhoff-6 in the details of the amino acid partitioning rationale, but the amino acid content of their bins broadly overlap (*5*). The Dayhoff-6 recoding partitions amino acids into six differently sized bins: one 5–amino acid bin, two 4–amino acid bins, two 3–amino acid bins and one 1–amino acid bin, based on how frequently they are expected to exchange with each other (i.e. their exchangeability rate). (**A**) the bins of the Dayhoff-6 recoding and the biochemical properties of the amino acids in each bin. (**B**) An exemplar amino acid dataset and its Dayhoff-6 recoded representation. Dayhoff-6 recoding is achieved by replacing, in a multiple sequence alignment, one letter amino acid codes with one letter codes representing the bins.

Every masking strategy induces a loss of information (*5, 19*). Yet, amino acid recoding of nucleotide data illustrates that, depending on the distribution of signal and noise, reducing the information content of a dataset can improve the ability of a dataset to resolve phylogenetic relationships (e.g. (*32, 33*)). Furthermore, it can be hypothesized that the impact of information loss should decrease as alignment size increases. Nonetheless, it is clear that information loss is potentially a problem when using recodings (*19*).

Fit is key to phylogenetic accuracy (*25, 34–36*) and it can be relative or absolute (*36*). Standard model fit tests, such as the Akaike Information Criterion (e.g. (*37*)), compare the relative fit of different models identifying the one that fits best relative to the others. However, a model of relative best-fit can be a poor descriptor of key properties (e.g. across-site compositional heterogeneity) of the data (*5, 36*). Goodness-of-fit tests (e.g. PPA) evaluate the absolute fit of models to the data (*36*), and can quantitatively express the extent to which we might be willing to trust results inferred using a model of relative best-fit (*5, 25, 36, 38–40*). The first attempt at testing the absolute fit of evolutionary models to amino acid and recoded data found that these tests achieve better (i.e. lower) PPA scores on recoded datasets, leading to the suggestion that recoding the data can improve model fit (*5*). Yet, this argument has been challenged because information loss can also lower PPA scores (*19, 41*).

We studied the effectiveness of recodings using simulated phylogenomic datasets and models that account for across-site compositional heterogeneity progressively better. We tested if the use of recodings can cause a deterioration of the phylogenetic signal, and whether their efficacy depends on alignment size. We tested whether recodings can induce the emergence of tree reconstruction artifacts and devised a new (recoding-based) strategy to test if clades are artifactual. We found that recodings substantially increase accuracy if the model used poorly fits the amino acid data, and the alignment is sufficiently long. If the model fits the amino acid data well, recoding did not improve fit nor accuracy, mildly decreasing support for key nodes instead. However, we find no evidence that the use of recodings might by itself cause the emergence of incorrect but statistically supported relationships (i.e. recodings are conservative). Our results are consistent with the hypothesis that recodings improve fit and consequently accuracy by reducing heterogeneity (*5*). Accordingly, recodings can only be beneficial when the model does not fit the amino acid data. We propose a guiding principle to decide when recodings can be useful, and confirm that many real datasets might benefit from recoding (*5*). We test our new approach to identify artifactual clades using the data of (*17*) to exemplify our approach while tackling one of the trickies nodes in the tree of life, the root of the animal phylogeny (*5–7, 12–19, 42, 43*). In so doing we found evidence that a placement of the ctenophores at the root of the animal tree is most likely artifactual.

## Results

### Simulated data and their heterogeneity

To avoid experimenter-induced biases, we did not simulate our own datasets. Instead we used the across-site compositional and rate heterogeneous alignments of (*18*), where the target (true) tree under which the data were generated assumed either that the ctenophores (Ctenophora-sister topology) or the sponges (Porifera-sister topology) are the sister of all the other animals. For each topology, this dataset includes 100 alignments of 30,000 sites and 97 taxa, see Methods. The alignments were subsampled to generate datasets of 1,000, 5,000, 10,000 and 30,000 sites and 20 taxa, as well as datasets with 30,000 sites and 4 or 10 taxa – see Supplementary Information (SI), Fig. S1, for the target trees. The alignments were recoded using the Dayhoff-6 amino acid recoding (described in Fig. 1). Unless otherwise stated, analyses were performed using the 30,000 sites and 20 taxa alignments, under both target topologies, see the flow chart of our experimental design in Fig. S2.

We used PPA of site-specific amino acid diversity (PPA-Div, (*44*)) to test whether the across-site compositional heterogeneity of the simulated amino acid alignments was comparable to that of real data. These results (reported in SI) confirmed that real and simulated data have comparable across-site compositional heterogeneity (Fig. S3), validating the use of the simulated alignments of (*18*) in our study.

### Dayhoff-6 improves accuracy if the model poorly describes the compositional heterogeneity of the amino acid data

We compared the success of amino acid and Dayhoff-6 recoded data under a series of models allowing for increasing amounts of across-site compositional heterogeneity. Throughout the study we define successful (see methods) an analysis correctly resolving the target topology with a minimal support, measured as its Posterior Probability (PP), of 0.5 (PP≥ 0.5). We distinguish two types of unsuccessful analyses, those that are incorrect (wrong tree inferred with PP≥ 0.5), and those that are unresolved (correct or incorrect tree inferred with PP< 0.5). We performed a sensitivity analysis where we increased the threshold for success to PP=0.95. This test did not affect our conclusions, see below and SI for details.

We started from a CAT-based model with 10 frequency categories (nCAT10 – to account for across-site compositional heterogeneity), which was combined with a Poisson process (F81, (*45*) that does not adjust for replacement rate heterogeneity) and a Gamma distribution (G) modeling across-sites rate heterogeneity. We progressively increased the number of frequency categories to 30, 60, 120 and 240 (to accommodate more across-site compositional heterogeneity - analyzing the data using nCAT30-F81+G, nCAT60-F81+G, nCAT120-F81+G and nCAT240-F81+G (hereafter referred to as nCAT10 to nCAT240). In addition, we tested the use of GTR+G (hereafter GTR), which cannot account for across-site compositional heterogeneity but models replacement rate heterogeneity, CAT-F81+G (hereafter CAT) which optimizes the number of categories used to accommodate across-site compositional heterogeneity during tree search, and CAT-GTR+G (hereafter CAT-GTR) which optimizes the number of categories used and replaces the Poisson process with a GTR matrix to model replacement rate heterogeneity.

Previous simulations using amino acid data, showed that it is easier to correctly infer Ctenophora-sister than Porifera-sister (*18*). While it is not a goal of this study to test why this might be the case, it has been suggested that this could be because Ctenophora-sister lies in the “Farris-zone” (*18*), having two long branched lineages (outgroups and Ctenophora – Fig. S1) on one side of the target branch and two short branched lineages (sponges and all the remaining animals) on the opposite side of the target branch (Fig. S1). Our analyses confirmed that Ctenophora-sister is easier to infer than Porifera-sister. Using amino acid data Ctenophora-sister is frequently inferred even when false, if the model has a poor fit (according to PPA) to the data. When Ctenophora-sister is the simulation target it is invariably inferred with a Success Rate (SR) close to 100%, irrespective of model and recoding (see Fig. 2a). On the contrary, Porifera-sister is only retrieved (and not 100% of the times) when true, with the model and coding strategy used having a strong impact on SR (Fig. 2a). More precisely, Dayhoff-6 increased SR when the model used poorly fit the amino acid data (Figs. 2a-c). To integrate the fact that SRs differ under Porifera– and Ctenophora–sister, for each model and coding scheme we calculated the Total Success Rate (TSR). For each model, this is the percentage of successful analyses, under both target topologies, when amino acids (TSR_AA_) or Dayhoff-6 (TSR_Day_) are used (Fig. 2b). We observed that the difference between PPA-Div_AA_ and PPA-Div_Day_ (hereafter referred to as *d*PPA-Div; measured as the difference in Z-score between PPAs performed using differently coded data – see Methods for a guide to interpret Z-score) correlates well (R^2^= 0.90) with the difference between TSR_Day_ and TSR_AA_ (hereafter *d*TSR – Fig. 2d). More precisely, as the evolutionary model better fits the amino acid data *d*TSR diminishes (Fig. 2d), becoming negative (meaning that TSR_AA_ was best) for models that fit the amino acid data well (CAT, nCAT240, and CAT-GTR – see Fig. 2c).

**Figure 2.**
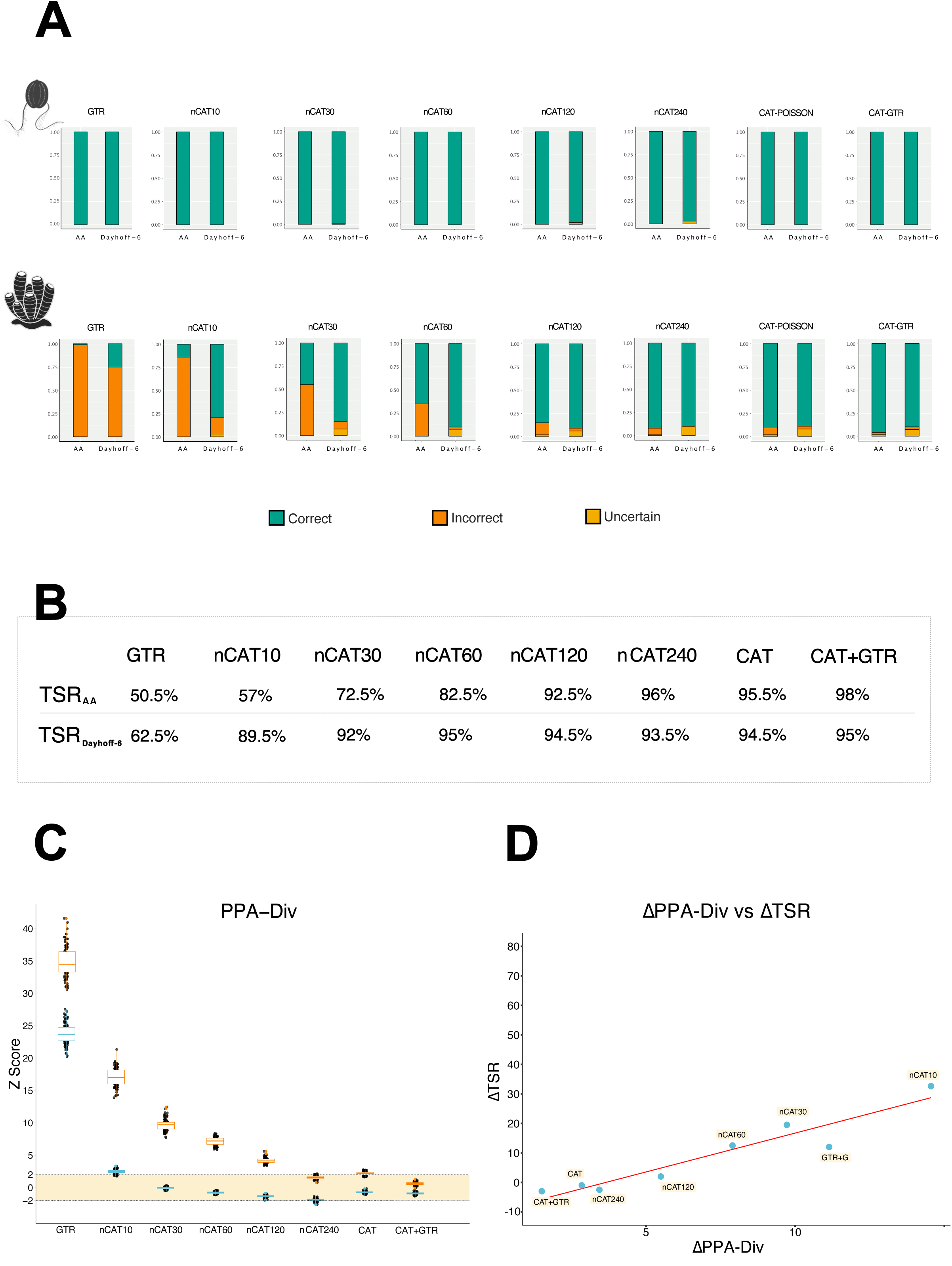
(**A**) Success rate of amino acids and Dayhoff-6 as models that can account for more across-sites compositional heterogeneity are used. (**B**) A table summarizing the Total Success Rate (TSR) for amino acids and Dayhoff-6 under each tested model. TSR is calculated (from the values in Fig. 2A) as the percentage of successful analyses (correct tree inferred with PP≥ 0.5) under both Porifera- and Ctenophora-sister – see also text. (**C**) Change in the fit of the model to the data (estimated using PPA-Div) as models that can account for more across-sites compositional heterogeneity are used. In Orange amino acid datasets; in Blue Dayhoff- 6 datasets. (**D**) Correlation between the difference in fit achieved by each considered model on the amino acid and Dayhoff-6 datasets (*d*PPA-Div), against the difference in TSR achieved before and after recoding (*d*TSR). All analyses were run in Phylobayes (Total number of cycles= 2,000; burnin= 1,000; subsampling frequency= 10 – see Sensitivity Analyses for a test of the validity of these settings).

The CAT-based model with the worst fit to the amino acid data was found to be nCAT10 (Average PPA-Div_AA_∼ 17; Fig. 2c). Dayhoff-6 data achieved much better PPA-Div scores for this model (Average PPA_Day_∼ 2.5), and nCAT10 emerged as the model for which recoding most strongly improved TSR (TSR_AA_= 57%; TSR_Day_= 89.5%; Figs. 2a,b,d – a 32.5% improvement). On the contrary, the model with the best fit to the amino acid data was CAT-GTR (Average PPA-Div_AA_= 0.6 – Fig. 2c). For this model PPA-Div_AA_ and PPA-Div_Day_ were comparable (Average PPA-Div_Day_∼ -0.9), suggesting that CAT-GTR fits both data types well. CAT-GTR emerged as the model with the worst relative performance under Dayhoff-6 (TSR_AA_= 98%; TSR_Day_= 95%; Fig. 2b – corresponding to a 3% reduction in TSR). Two more models were found to fit the amino acid data well: nCAT240 (Average PPA-Div_AA_ ∼1.5) and CAT (Average PPA-Div_AA_ ∼2). As in the case of CAT-GTR, PPA-Div_AA_ and PPA-Div_Day_ were comparable for these models, suggesting a good fit to both data types, and as in the case of CAT-GTR, the performance of these models did not improve upon recoding (Figs. 2a and 2d). Importantly, the reduced TSR_Day_ observed for modes that fit the amino acid data, did not result from the recovery of incorrect trees, but by an increment in the rate of unresolved outcomes. These results suggest that when the model is already fitting the amino acid data well, we suffer the cost of recoding (information loss), without any benefit (no violation left to palliate). The other CAT-based models tested (nCAT30-nCAT120) had PPA-Div_AA_ falling within the extremes defined by nCAT10 and CAT (Fig. 2c) and showed intermediate increments in TSR (Figs. 2b and d), which become progressively less marked as PPA-Div_AA_ improved. Finally, GTR performed worse than any of the CAT-based models on both data types, with PPA-Div_AA_ and PPA-Div_Day_ suggesting that this model fit very poorly both types of data (Average PPA- Div_AA_= 43.6; Average PPA-Div_Day_= 30.2). Consistently with this observation and with the fact that PPA-Div_Day_ potentially indicates a marginally better fit of this model on Dayhoff-6 data, recoding improved the TSR of GTR by 12%, but in a context where, under Porifera-sister, this model was still three times more likely to recover an artifactual Ctenophora-sister topology rather than the target tree.

### Information loss cannot explain results obtained using Dayhoff-6

It has been suggested that the topological changes and reduced PPA scores observed when amino acid alignments are recoded might be artifacts caused by signal loss (*19*). We formally tested this hypothesis using a jackknife approach. To achieve our goal, we stripped sites from our amino acid datasets to simulate the effect of a loss of information comparable to that that Dayhoff-6 would impose if every substitution in the amino acid alignment was phylogenetically informative. This is a worst-case scenario as no dataset is homoplasy-free, and hence the true amount of information that Dayhoff-6 can strip from the data should be lower than the amount we estimate. Accordingly, the results of our test should be conservative and bias our results against Dayhoff-6. For this experiment, datasets were analyzed using nCAT10. This model was chosen because (see Fig.2) it most strongly highlights differences between results obtained using Dayhoff-6 and amino acids.

The amino acid datasets were found to have an average tree length of 100,914.18 steps, while the corresponding Dayhoff-6 datasets were found to have an average tree length of 49,683.18 steps. If we assume each substitution to be phylogenetically informative, we can use tree length as a measure of the phylogenetic signal in the data, and we can estimate that Dayhoff-6 would reduce the signal in the data by ∼49%. We thus jackknifed the amino acids datasets to remove ∼49% of their sites. The average tree length of the jackknifed datasets was 51,179.6 steps, which is comparable (even if not identical) to the average tree length calculated for the Dayhoff-6 datasets (Fig. 3a), indicating that removing 49% of the sites from our amino acid datasets generated jackknifed datasets with a number of substitutions comparable to that observed in the Dayhoff-6 datasets. We calculated PPA-Div scores for the jackknifed datasets and compared these values against those achieved by full-length amino acid and Dayhoff-6 datasets. PPA-Div scores for the Jackknifed datasets were significantly worse (Average PPA-Div_Jack_= 12; SD= 0.99) than those of the Dayhoff-6 datasets (Average PPA-Div_Day_= 2.47; SD=0.33) and remained comparable to those inferred from full-length amino acid datasets (Average PPA-Div_AA_= 17.32 SD= 1.4; Fig. 3c). The comparison of PPA- Div_AA_ and PPA-Div_Jack_ shows that reducing information indeed mildly reduced PPA-Div scores. However, the comparison of PPA-Div_Jack_ and PPA-Div_Day_ scores demonstrates that a loss of signal comparable to that imposed by the application of Dayhoff-6 cannot explain PPA-Div_Day_ scores, which are significantly better (the distributions do not overlap) than both PPA-Div_AA_ and PPA-Div_Jack_ (Fig. 3c). Furthermore, the TSR of the jackknifed datasets (TSR_jack_= 60.5%) is comparable to that achieved by the full-length amino acid data (TSR_AA_=57% – Fig. S4a) and both are much worse than TSR_Day_ (TSR_Day_= 89.5%; Fig. 2b and S4a). Significantly better TSR_Day_ values cannot be caused by a deterioration of the signal in the data, rejecting the hypothesis that results of Dayhoff-6 are driven by signal loss.

**Figure 3.**
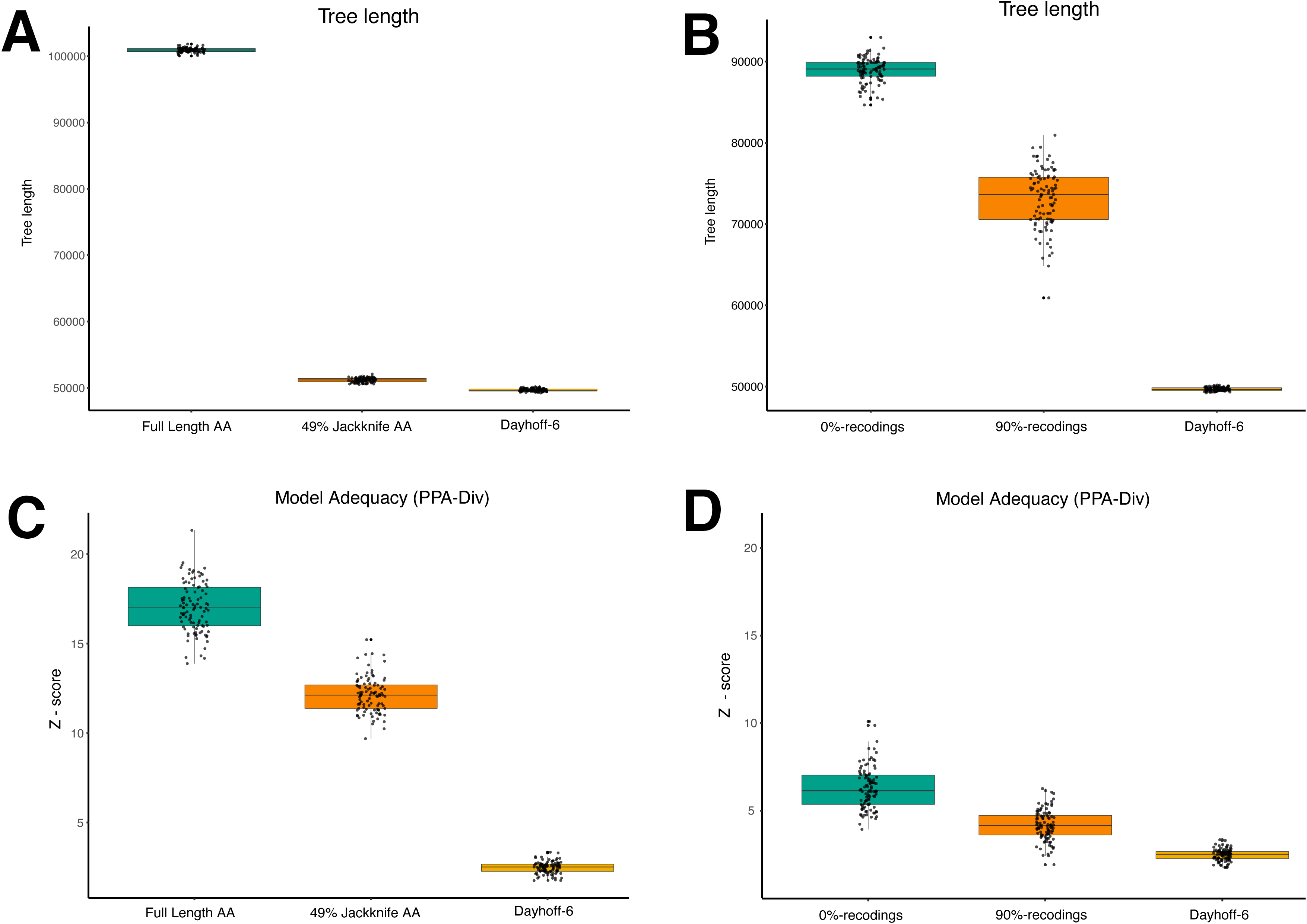
(**A**) Boxplot representing the distribution of tree lengths for the full length amino acid datasets, the 49% Jackknife amino acid dataset, and the Dayhoff-6 datasets. (**B**) Boxplot representing the distribution of tree lengths for 0%-recoded, 90%-recoded, and Dayhoff-6 recoded dataset. (**C**) PPA-Div scores for the full length amino acid datasets, the 49% Jackknifed amino acid datasets, and the Dayhoff-6 datasets. (**D**) Comparison of PPA-Div scores for amino acid datasets, 0%-recoded, 90%-recoded, and Dayhoff-6 recoded datasets. The figure indicates that PPA-Div scores of Dayhoff-6 datasets are significantly lower (i.e. better) than PPA-Div scores from jackknifed datasets – panel C, and 0%-recodings – panel D, as the distributions do not overlap. PPA-Div refers to analyses performed under nCAT10 in Phylobayes (Total number of cycles= 2,000; burnin= 1,000; subsampling frequency= 10 – see Sensitivity Analyses for a test of the validity of these settings).

### Random recodings confirm the efficacy of Dayhoff-6

The efficacy of recodings can be investigated by comparing their results to those of randomly generated ones. The latter are recodings where amino acids are randomly reassigned across the bins of an exchangeability-informed (parent) recoding (*19, 31*) – hereafter simply referred to as the “parent”.

The theoretical rationale that underpinned the development and application of recodings is grounded on the prediction that there should be more within-bin substitutions than across-bin substitutions (see introduction and Fig.1). If this theoretical prediction is to hold, randomly generated recodings should be expected to mask less substitutions than their parent, and phylogenies inferred from randomly recoded datasets should have higher tree length. In addition, random recodings should not be expected to reduce compositional heterogeneity to the same level of their parent. This is because the masked substitutions, being randomly selected, would not necessarily be between amino acids with similar properties (*19*). However, it is important to note that random recodings do not randomize the information in the data, they randomly select the subset of substitutions to mask (see Fig. 4). Accordingly, random recodings are Jackknife-like procedures and their application is expected to return non-random trees that will be accurate and well supported if the original amino acid alignment was informative. Nonetheless, the application of random recodings should not be expected to improve accuracy to the same extent of the parent recoding (*19*). Finally, we need to consider that there is only a limited number of ways in which 20 amino acids can be partitioned across six bins, and random recodings can be expected to retain residual amino acid similarity to their parent.

**Figure 4.**
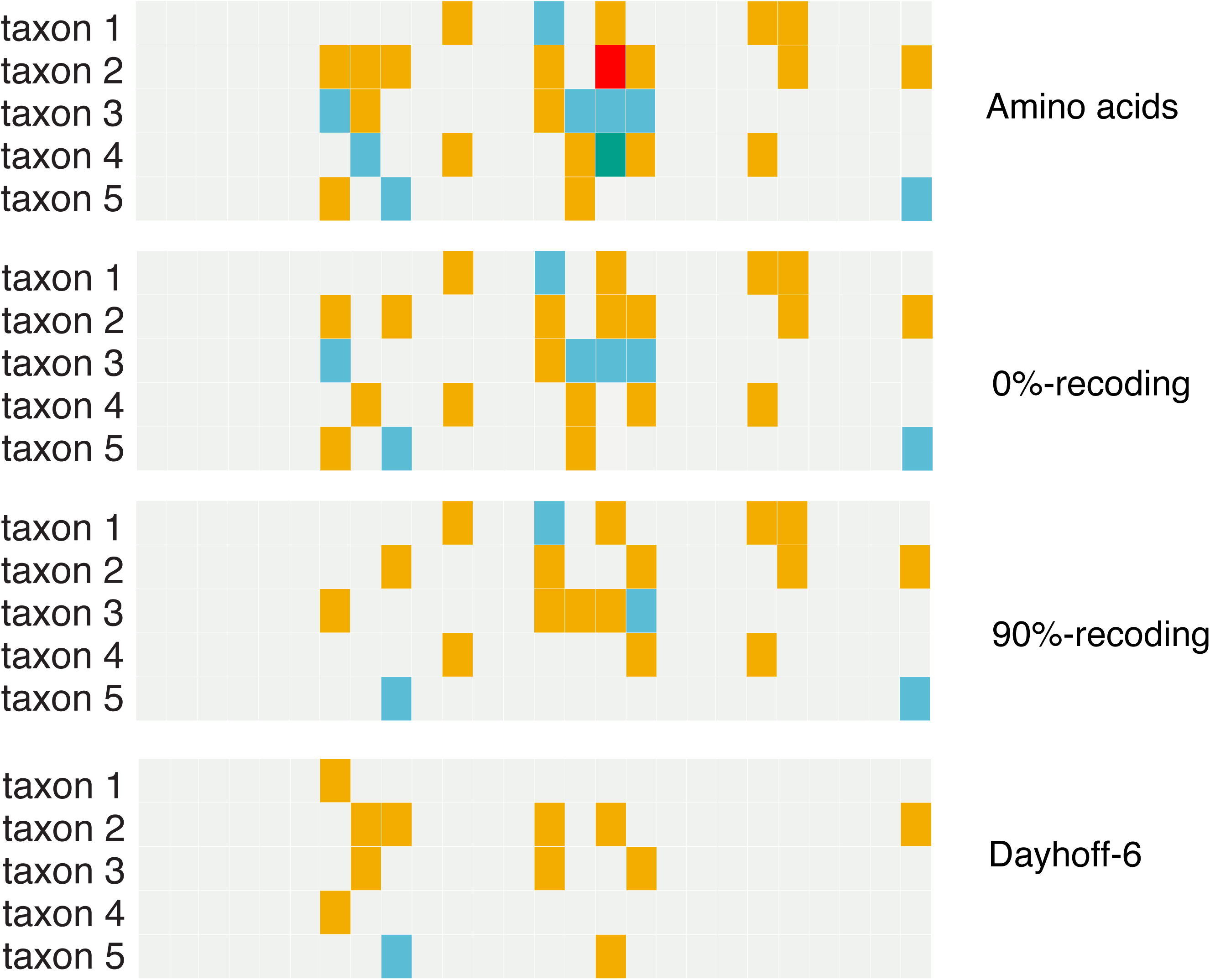
A graphical representation of the similarities between the 0%-, 90%- and Dayhoff-6 recodings of the same amino acid dataset. The alignment represents the first 26 sites of a randomly chosen simulated dataset. The figure shows that there is a clear similarity between the various coding strategies, i.e. random recodings do not scramble the signal in the data; they subsample it. The figure shows that Dayhoff-6 masks more substitutions than the random recodings, indicating that within-Dayhoff-6-bin substitutions are more abundant than across- Dayhoff-6-bin substitutions. Furthermore, the figure shows that as recodings become more similar to Dayhoff-6 they mask more substitutions (see also Fig. 3 and main text). Note that, in random recodings, amino acids are randomly reassigned to bins, so a common naming system for the bins of different recodings cannot be achieved. Hence in the figure we only use colors. If for a site, all species have the same state (either the same amino acid in the case of the amino acid alignment, or if all amino acids observed at that site fall into the same bin in the case of the recodings), we use a single color to represent the site. If two states are present (either two amino acids or the observed amino acids fall in two different bins) we use two colors. If three amino acids are present or the observed amino acids fall in three bins, we use three colors. The coloring of each column, therefore, simply represents the number of amino acids or bins observed at each position. Across sites, the same color was used for the most common state (irrespective of the bin name for that state), the second most common state, and so on.

If the rationale underpinning the use of recodings holds and recodings work as intended, we can thus make three predictions. (I) Phylogenies inferred from randomly recoded datasets will have a higher treelength than those inferred from datasets recoded using the parent recoding. In addition, we expect to observe that the length of trees inferred from randomly recoded datasets will decrease as the residual amino acid similarity between the random recodings and the parent recoding increases. (II) PPA-Div scores of randomly recoded datasets will be worse (higher Z-scores) than those of datasets recoded using the parent recoding. However, we expect Z-scores of randomly recoded datasets to progressively improve as the residual amino acid similarity between the random recodings and their parent increases. (III) The analysis of randomly recoded datasets can be expected to return non-random trees, but these trees will be less accurate (lower TSR) than those inferred using the parent recoding. However, the TSR of randomly recoded datasets is expected to improve as the similarity between randomly generated recodings and their parent increases.

We generated 1,000 random recodings (see Methods) based on the Dayhoff-6 binning scheme, finding that they retain on average 36.5% similarity to Dayhoff-6 (range 0% – 80%; see Supplementary Table 1 - https://bitbucket.org/bzxdp/giacomelli_et_al_2022_recodings/src/master/). We then identified 100 recodings of 0% and 90% residual amino acid similarity to Dayhoff-6 (hereafter 0%- recodings and 90%-recodings) using a process where random recodings were generated, compared to their parent, and retained only if they had the desired residual amino acid similarity to Dayhoff-6 (see Methods). We used the 0%- and 90%-recodings to test our three predictions.

Our results indicate that the extent to which randomly generated recodings mask substitutions and improve PPA-Div and TSR scores depends on their similarity to Dayhoff-6. 0%-recodings had an average tree length of 88,046.08 steps and 90%-recodings of 72,390.12 steps, indicating that Dayhoff-6 silence ∼39% and ∼23% more substitutions than 0%- and 90%-recodings (Fig. 3b). PPA-Div scores obtained from Dayhoff-6 (Average PPA-Div_Day_∼ 2.5; SD= 0.33) are significantly better (the distributions do not overlap) than those obtained using 0%-recodings (Average PPA-Div_0%_∼ 6; SD= 1.2), while the distribution of PPA-Div scores obtained using 90%-recodings (Average PPA-Div_90%_ ∼ 4; SD= 0.87) overlap with those of both the 0%-recodings and Dayhoff-6 (Fig. 3d). Finally, we found that the level to which randomly generated recodings improve accuracy (TSR_0%_= 81%; TSR_90%_= 83.5%) depends on their similarity to Dayhoff-6 (TSR_Day_= 89.5) – Fig. S4b. More precisely, when the target is Ctenophora-sister SR= 100% irrespective of recoding strategy (Figure S4b), while when the target is Porifera-sister SR_0%_= 62%, SR_90%_= 67%, SR_Day_= 79%.

### The success of Dayhoff-6 increases with the length of the alignment

Recodings mask substitutions and therefore induce a loss of signal. While this loss of signal does not negatively affect results obtained using our 30,000 sites datasets (see above), we hypothesize that the efficacy of recodings should be alignment-length dependent. To test this hypothesis, we analyzed alignments with 20 taxa (Fig. S1) and 1,000, 5,000, 10,000 and 30,000 sites, simulated under both Ctenophora- and Porifera-sister (see above and methods) using nCAT10.

We found that with 1,000 site alignments, amino acids perform better than recodings TSR_AA_= 26% and TSR_DAY_= 11% (Fig. 5a and Fig. S5). Amino acids perform marginally better than Dayhoff-6 also with alignments of 5,000 sites (TSR_AA_= 59.5% and TSR_Day_= 58.5% – Fig. S5). However, with alignments of 10,000 sites the overall performance of Dayhoff -6 is better than that of amino acid data (TSR_AA_= 63% and TSR_Day_=74%; Fig. S5). However, with the 10,000 site alignments, Dayhoff-6 does better than amino acids under Porifera-sister (SR_AA_= 26%; SR_AA_= 54%) while amino acids are still slightly better than Dayhoff-6 under Ctenophora- sister (SR_AA_= 100%; SR_Day_= 94%). Finally, with 30,000 site alignments Dayhoff-6 outperforms amino acids (TSR_AA_= 57% and TSR_Day_= 89.5%; Fig. 5b and S5), with Porifera-sister being correctly inferred only 14% of the times using amino acids and 79% of the times under Dayhoff- 6, when this clade is true.

**Figure 5.**
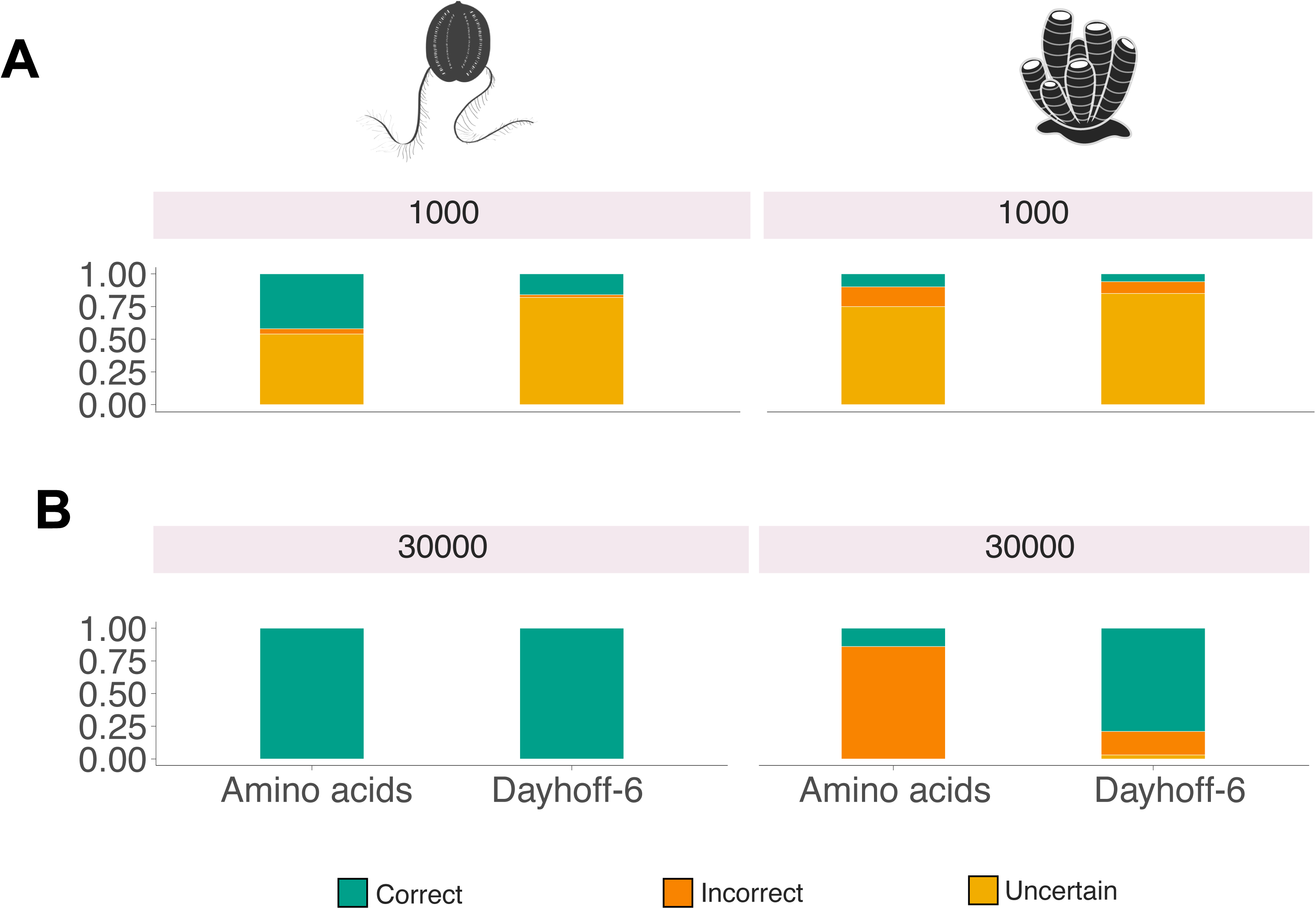
(**A**) Success rate of 1,000 sites alignments recoded as amino acid and Dayhoff -6 datasets under Ctenophora-sister and Porifera-sister. (**B**) Success rate of 30,000 sites alignments recoded as amino acid and Dayhoff-6 datasets under Ctenophora-sister and Porifera-sister. All analyses were performed under nCAT10. The figure shows that with 1,000 site alignments amino acids perform better than Dayhoff-6, but with 30,000 site alignments Dayhoff-6 performs better than amino acids. See Figure S5 for the performance of amino acids and Dayhoff-6 with 5,000 and 10,000 site alignments. All analyses were run in Phylobayes (Total number of cycles= 2,000; burnin= 1,000; subsampling frequency= 10 – see Sensitivity Analyses for a test of the validity of these settings).

### Sensitivity analyses

We tested the effect of our most important experimental choices. We investigated whether the use of Bayesian inference instead of Maximum Likelihood could have affected our results, we studied the effect of changing the number of cycles and runs in our Bayesian analyses, the use of alternative recoding protocols (SR6 and KGB6 – Fig. 1), and the use of a more stringent threshold to define a successful analysis (PP= 0.95). We investigated whether taxon sampling density affected our results (subsampling our 30,000 sites datasets to include 10 or 4 taxa), and we tested the effect of combining recodings with an overparameterized CAT-based model that used 544 frequency categories (nCAT544), twice the number that CAT optimally infers on average for our simulated data. The results of these analyses are presented in detail in SI (particularly in Figs. S6 to S11) and show that our results were not affected by our methodological choices. In particular, we found no evidence that combining recodings with an overparameterized model might result in the inference of artifactual trees, as the TSR of nCAT544 remains excellent, irrespective of the coding strategy used (TSR_AA_= 96.5%; TSR_Day_= 94%).

### Application of randomly generated recodings to real datasets can be used to identify artifactual clades

We applied the knowledge gained from our simulations to a real dataset. We used the dataset chosen by (*19*), Whelan-strict (*17*). We analyzed this dataset as an amino acid alignment, as well as using Dayhoff-6, 0%- and 90%- recodings using nCAT60, as in (*19*). We estimated PPA_AA_, PPA_0%_, PPA_90%_, and PPA_Day_ scores, as well as support values for both Porifera- and Ctenophora-sister under all coding strategies.

Whelan-strict is expected to be affected by both across-site and across-lineage compositional heterogeneity, as it is generally the case for real datasets. We thus used both PPA-Div and PPA-Max (which tests how well a model describes across-lineage compositional heterogeneity (*5*)) to test the fit of nCAT60 to the data. PPA-Div_AA_ indicates that nCAT60 describes the across-site compositional heterogeneity of Whelan-Strict poorly (PPA-Div_AA_∼19). PPA-Max indicates that nCAT60 describes relatively well the across-lineage compositional heterogeneity of Whelan-strict (PPA-Max_AA_∼ 2.65), and this form of heterogeneity is thus not expected to represent a major problem for this dataset under nCAT60. The PPA-Div score obtained for Whelan-Strict under nCAT60 is comparable to those obtained (for our simulated data) using nCAT10 – the model for which Dayhoff-6 improved TSR the most (Fig. 2b). Furthermore, PPA-Div_Day_ indicates that while nCAT60 describes the across-site compositional heterogeneity of the amino acid data poorly, it fits well the Dayhoff-6 data (PPA-Div_Day_ = -0.9). PPA-Max_Day_= 1.91 indicates that nCAT60 also adequately describes the across-lineage compositional heterogeneity of the Dayhoff-6 data. As expected, values for PPA-Div_0%_ and PPA-Div_90%_ achieved intermediate values (PPA-Div_0%_ ∼5.4; PPA-Div_90%_ ∼2.7). Based on our simulations, these results are interpreted to suggest that there is scope for recodings to improve the accuracy of nCAT60 with this dataset.

We found that support for Ctenophora-sister was maximal when the data were analyzed as amino acids (Fig. 6) and progressively decreased, reaching minimal support when the data were Dayhoff-6 recoded (Fig. 6). Support for Porifera-sister followed the opposite trend, achieving maximal support under Dayhoff-6 (Fig. 6). We plotted the average support values observed under nCAT10 (the model that in our simulations achieved PPA-Div_AA_ scores comparable to those obtained using nCAT60 for Whelan-Strict) when both Porifera- and Ctenophora-sister are true (Fig. 6). Changes in support values observed for Porifera-sister are comparable to those observed, using simulated data, when Porifera-sister was true. Differently, the pattern observed for Ctenophora-sister disagrees with what observed in simulations where Ctenophora-sister was true (Fig. 6). The pattern of change in support observed for Porifera-sister is in agreement with what (*19*) suggested we should observe, if recodings were effective at improving phylogenetic accuracy (and our results suggest that they are) and sponges were the sister of all the other animals. Differently, the pattern of change in support values observed for Ctenophora-sister is consistent with what should be observed if Ctenophora-sister was a tree reconstruction artifact (*19*).

**Figure 6.**
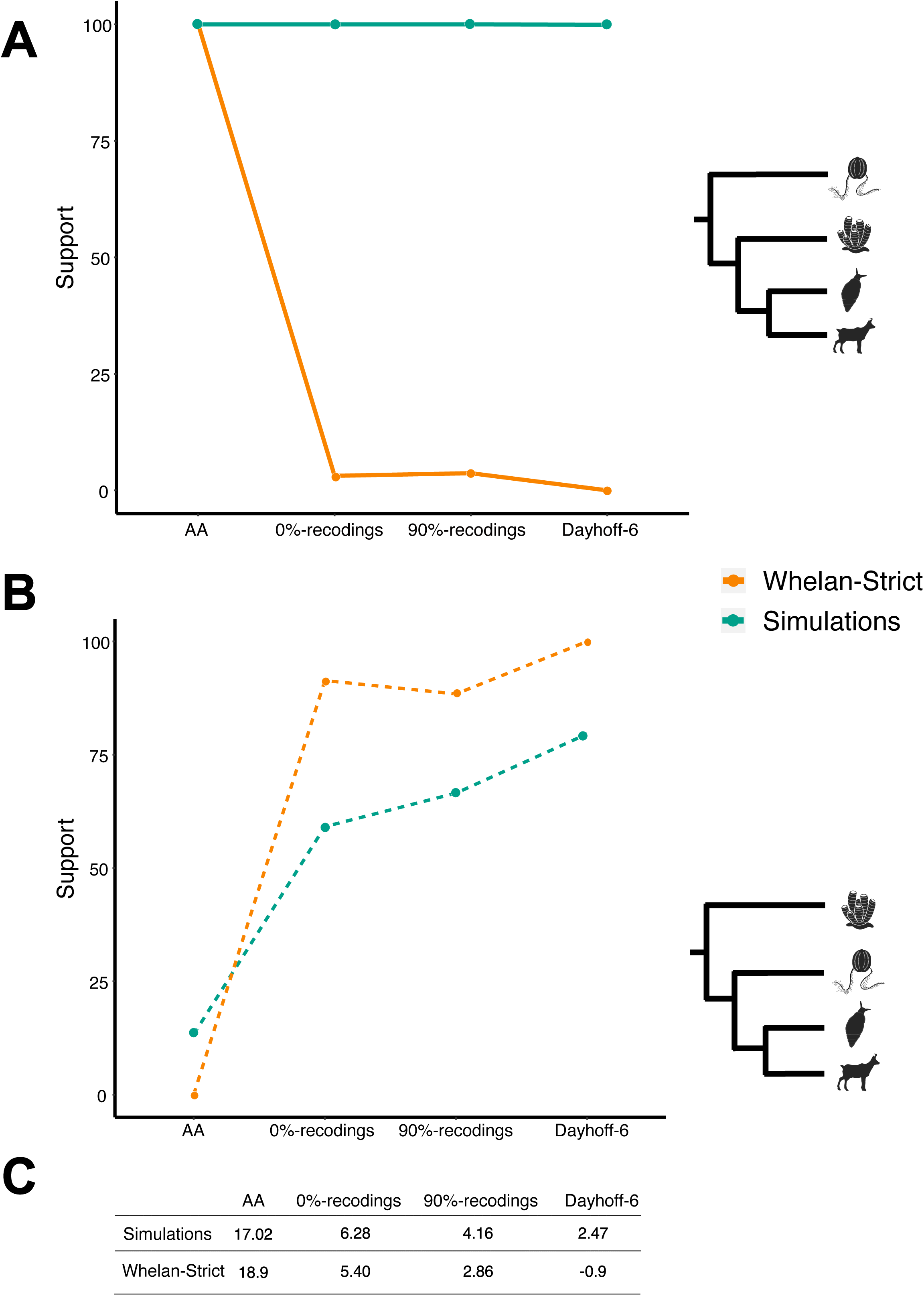
Analyses of Whelan-strict. (**A**) In Orange: Changes in support for Ctenophora-sister as the data were coded as amino acids, 0%-, 90%- and Dayhoff-6 data. Analyses were performed under nCAT60. In Green: Support values obtained in our simulations when Ctenophora-sister is true. The simulated data were analyzed under nCAT10 (see main text and point C below for the justification for the use of this model). (**B**) In Orange: Changes in support for Porifera-sister as the data were coded as amino acids, 0%-, 90%-, and Dayhoff-6 data. Analyses were performed under nCAT60. In Green: Support values obtained in our simulations when Porifera-sister is true (under nCAT10). (**C**) Top: PPA-Div scores for the simulated datasets (under nCAT10) when the data are coded as amino acids, 0%-, 90%- and Dayhoff-6 data. Bottom: PPA-Div scores for Whelan-strict, under nCAT60, when the data are coded as amino acids, 0%-, 90%- and Dayhoff-6 data. Note that analyses were repeated using the complete set of one hundred 0%-recodings and 90%-recodings hence support values and PPA-Div scores for these recodings are average values across 100 analyses.

## Discussion

### The use of recodings is justified

The use of recodings have been theoretically justified based on their predicted ability to mask substitutions that are *a-priori* expected to be more likely to contribute to the compositional heterogeneity of a data (the within-bin substitutions; e.g. (*5*)). However, (*19*) correctly pointed out that there is no published empirical evidence to suggest that the use of recodings is justified. Our random recoding experiments in conjunction with our amino acid jackknife experiment show that: (1) within-Dayhoff-6-bin substitutions are more abundant than across- Dayhoff-6-bin substitutions as Dayhoff-6 remove more substitutions than random recodings; (2) within-bin substitutions contribute more to the compositional heterogeneity of amino acid datasets than across-bin substitutions as Dayhoff-6 improves PPA-Div scores more than random recodings; (3) Dayhoff-6 is effective at improving fit as Dayhoff-6 PPA-DIV scores are significantly lower than those obtained using jackknifed and randomly recoded datasets; (4) Dayhoff-6 improves phylogenetic accuracy as it achieve higher TSR than randomly generated recodings and jackknifed amino acid datasets. Overall, these results indicate that the theory underpinning the use of recodings holds. In addition, these results are consistent with the hypothesis that recodings improve accuracy because, by reducing compositional heterogeneity, they can improve the fit of the model to the data (*5*). In particular, an improved TSR cannot be explained by a deteriorating signal, and this result is the strongest evidence for the utility of recodings when tackling tricky nodes. The tests of (*19*) failed to reach these conclusions because it used only four random recodings and did not consider that they retained spurious amino acid similarity (10% to 40%) to their parent (SR6).

Notably, even 0%-recodings outperformed amino acids (compare Figs 2, 3 and S4). It might seem counterintuitive that 0%-recodings can achieve greater TSR (TSR_0%_= 81%) than amino acids (TSR_AA_∼ 57%) and it is not the scope of this paper to investigate this phenomenon. However, it should not surprise that 0%-recodings can infer good quality trees given that randomly recoded datasets are not randomized, they are jackknifed.

### Recodings improve accuracy if the model used is not a good descriptor of the compositional heterogeneity of the amino acid data and the alignment is sufficiently long

The relative performance of recoded and amino acid datasets had not been previously investigated using genome scale datasets and models that accommodate compositional heterogeneity progressively better. We show that when phylogenomic-scale alignments are used recodings improve accuracy if the model used had a poor fit to the amino acid data. This might be erroneously interpreted to mean that recodings are not useful if the data are analyzed under the best fit model. That is not the case, as model-fit tests compare the relative fit of models against each other (*36*), and it has been shown (e.g. (*5*) and this paper) that relative best fit models can fail absolute goodness-of-fit tests. When these conditions apply (in our study these are the cases of GTR, nCAT10, nCAT30, nCAT60 and nCAT120) phylogenetic accuracy can be improved recoding the data (Fig. 2). On the other hand, when the model fits well the amino acid data (in our study nCAT240, CAT and CAT-GTR) recoding does not improve accuracy. These observations are consistent with the hypothesis that recodings improve accuracy when they allow models to achieve a better fit to the data (*5*), and can thus only be useful when the model does not already fit the amino acid data well.

Recoding data masks substitutions and induces a loss of information. A recent simulation study (*41*) used alignments of 1,000 to 5,000 sites and concluded that recodings have an invariably deleterious effect on accuracy, challenging more than 100 papers that implemented such approaches. Our results similarly found that with 1,000 and 5,000 sites alignments, amino acids perform better than recodings. However, our results also show that the performance of the recoded data improves as alignment size increases, and with phylogenomic-scale (30,000 site alignments) datasets, Dayhoff-6 clearly outperforms amino acids (when the model used did not fit the amino acid data well – which was the case also in (*41*)). Our results reject the conclusion of (*41*), but also indicate, as we predicted, that the efficacy of recodings is alignment size dependent, and their application should be limited to phylogenomic analyses such as those in (*5, 6, 8, 10, 16, 17*) and in this study.

### A criterion to decide when to use recodings

Recodings improve accuracy only when their application allows improving fit, and a criterion is needed to decide when they should be used (*7, 41*). We show that the difference in the goodness-of-fit of the model to the amino acid and recoded data (*d*PPA) correlates strongly (R^2^=0.90) with the difference between the total success rate of recoded and amino acid data (*d*TSR – Fig. 2). Although it is not clear, on formal grounds, whether PPA-Div_AA_ and PPA- Div_Day_ values should be comparable since they are summary statistics computed on data defined on different state spaces, the trend in Figs. 2d suggests that these values are nevertheless able to predict if phylogenetic success can be improved using recodings. It should be noted that even if we did not directly compare these values, high PPA-Div scores would still indicate that the model has a lower probability of accurately resolving a phylogeny, compared to a model with a low PPA-Div score. Accordingly, an independent reading of PPA- Div_AA_ and PPA-Div_Day_ values (see Table 1), would still allow deciding whether a dataset is best analyzed as an amino acid or a Dayhoff-6 alignment. For example, following the indications in Table 1, one would conclude that under GTR there is a low chance of correctly resolving the target topology irrespective of recoding. Under nCAT10 we would conclude that there is a high probability of correctly resolving the target topology using the recoded data but not the amino acid data, and under CAT-GTR, both data types have a high chance of correctly recovering the target topology.

**Table 1.**
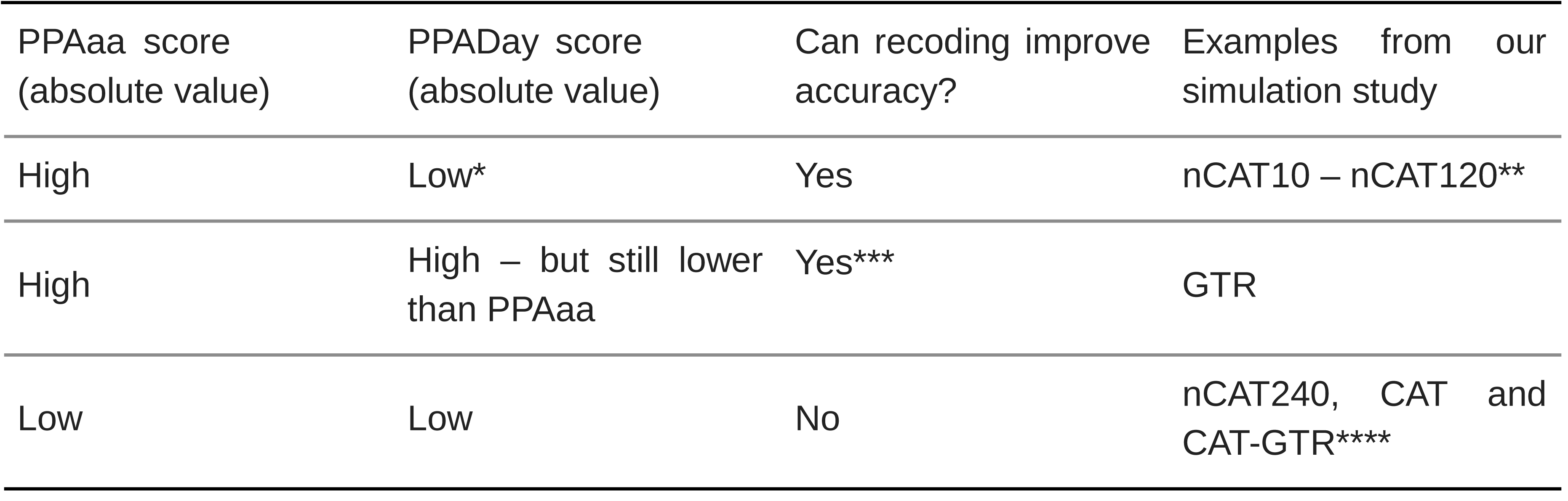
A simple guide to the application of exchangeability-informed recodings. PPA scores are here intended as absolute values. PPA scores approximately included between 2 and -2 indicate that the model is adequate (Methods for more details). *As a rule of thumb PPA score with an absolute value lower than 2 is considered low. If a model has a PPA_Day_ score with an absolute value higher than two, it might be worth considering the use of a model that can accommodate greater amounts of compositional heterogeneity if available. If such a model is not available our simulations suggest that even with a PPA_Day_ score higher than 2, as in the case of nCAT10, an substantial improvement in success rate can be observed, if PPA_AA_ indicated a poor or very poor fit to the data (see methods for details). **Note that as PPA_AA_ and PPA_Day_ become closer in value (i.e. as models that can accommodate greater amount of compositional heterogeneity are used and the fit of the model to the amino acid data progressively improves) the positive effect of using an exchangeability- informed recoding decreases (as there are less violations to palliate – see Fig. 2). ***When PPA_AA_ and PPA_Day_ scores are high, but PPA_Day_ is lower than PPA_AA_, an improvement in success rate can still be expected but in the context of an analysis where the chances of inferring an incorrect tree can still be high (as in the case of GTR). ****When both PPA_AA_ and PPA_Day_ scores are low (absolute values equal or lower than 2), the model fits both data types well and the use of recodings cannot be expected to improve fit nor accuracy, as there are no violations to palliate.

| PPAaa score<br>(absolute value) | PPADay score<br>(absolute value) | Can recoding improve<br>accuracy? | Examples from our<br>simulation study |
| --- | --- | --- | --- |
| High | Low* | Yes | nCAT10 – nCAT120** |
| High | High – but still lower<br>than PPAaa | Yes*** | GTR |
| Low | Low | No | nCAT240, CAT and<br>CAT-GTR**** |

A fundamental aspect of our conclusions is that recodings do not work best with a specific model: that is, our results do not suggest that Dayhoff-6 is best combined with nCAT10. Rather, our results suggest that the effectiveness of recodings depends on their ability to improve the fit of the model to the data. Simulated data lack the complexity of real data and PPA_AA_ of real data (this study and (*5*)), indicate that models such as nCAT60 and CAT-GTR can fail to fit real amino acid datasets. Accordingly, combining compositionally heterogeneous models with data recodings have the potential to improve fit and phylogenetic accuracy of real datasets.

### Ctenophora-sister is likely to be a tree reconstruction artifact

Results of posterior predictive analyses (PPA_AA_) of real datasets assembled to resolve the relationships at the root of the animal tree suggest that these datasets could benefit from recoding (Fig. 6; (*5*)). We agree with the suggestion of (*19*) that, if recodings improve accuracy, as indicated by the results of our simulation study, we should be able to use recodings to test if a clade is artifactual or not, as support for artifactual clades should be maximal using amino acid data and minimal using recoded data. Support for true clades should follow the opposite trend, as long as the conditions in which the application of recodings can be useful are met (see above). We should add to the suggestion of (*19*) that if random recodings were also used, artifactual clades should be expected to lose support as random recodings more similar to their parent are used, while support for true clades should follow the opposite trend. We tested our approach using Whelan-strict (*17*) under nCAT60, and found that PPA_AA_ and PPA_Day_ scores suggest that the use of recodings could improve accuracy for this dataset. When plotting support values for Porifera- and Ctenophora-sister using different coding strategies (amino acids, 0%-, 90%- and Dayhoff-6 recoding), we found that the trend observed for Porifera sister was that expected for true clades, while the trend observed for Ctenophora-sister was that expected for tree reconstruction artifacts (Fig. 6). The predictive power of these trends could be confirmed using our simulated data. We thus conclude that our test suggests that Ctenophora-sister is likely to represent a tree reconstruction artifact, and that his new approach can be broadly used to test whether specific resolutions of tricky nodes might represent tree reconstruction artifacts.

The debate on the root of the animal tree has seen a diversity of analyses and data. Sponges as the sister of all the other animals is favored by morphology and gene content data (*46–48*). Amino acid datasets analyzed using CAT and CAT-GTR tend to switch between Porifera– (*14*) and Ctenophora–sister (*49*) depending on outgroup sampling. In these experiments Porifera-sister is frequently (but not always) supported when close outgroups, which should reduce attraction artifacts (*5, 7, 14–17, 50*), are used. A common feature of all phylogenomic analyses of early animal relationships is that the models used to analyze the amino acid superalignments they use fail to a smaller (with CAT-based models) or larger (with GTR) extent to fit the data (*5*). When the relevant datasets are recoded, CAT-based models (but not GTR) seem to achieve a good fit ((*5*) and above). These are the conditions in which we expect that the use of recodings can improve phylogenetic accuracy, and the analysis of recoded data using CAT-based models such as CAT-GTR (or nCAT60 – see above), almost invariably find outgroup-independent support for Porifera-sister – but see (*51*) for an exception. Under the interpretative framework of (*5*), recodings improve model fit and should mitigate attraction artifacts. Accordingly, the broadening of support for Porifera-sister in analyses of recoded datasets was interpreted by (*5*) as an indication that Ctenophora-sister is a tree reconstruction artifact. Differently, (*19*) suggested that the results of (*5*) and of other studies supporting Porifera-sister are either affected by overparameterization-induced artifacts caused by the use of CAT, by biases caused by a deterioration of signal caused by the use of recodings, or, by extension, by a combination of these two factors.

Overparameterization in Bayesian analysis has been studied (*25, 52, 53*). These studies concluded that overparameterization can reduce support values (decreases precision), but does not cause the emergence of tree reconstruction artifacts, as topological errors disappear as alignment length increases as first noted by (*25*) and confirmed by (*53*). We performed our own simulations using a highly overparameterized nCAT544 model, and found, in agreement with (*25, 53*) that it only mildly reduced support values. Similarly, our simulations did not find any evidence that the use of recodings can cause artifacts associated with signal deterioration, not even when Dayhoff-6 is combined with nCAT544. Even in instances where the use of recodings is inappropriate (i.e. when the model already fits the amino acid data), their application only led to mild reductions in support value, indicating that their use is conservative. In particular, we did not find any evidence that Porifera-sister might be inferred as a tree reconstruction artifact, when Ctenophora-sister is the target (*19*).

It has been suggested that we should be skeptical of Porifera-sister because it is found only under a narrow range of models that do not fit the data better than models supporting Ctenophora-sister (*19*). However, the size of the model space supporting Ctenophora-sister is not as broad as suggested by (*19*) because WAG, LG, JTT, etc. (*54–56*), are just alternative parameterizations of the same model: GTR. If one were to consider alternative parametrizations of CAT as if they were independent models, the range of models under which Porifera-sister is supported will also expand. Furthermore, none of these alternatively parametrized GTRs fits as well as CAT or CAT-GTR (which also frequently support Porifera- sister, e.g. (*14, 57*), contrary to what seems to be implied by (*19*)). Indeed, (*19*) only showed that nCAT60 (another CAT-based across-site compositionally heterogeneous model) could not be distinguished in their Bayesian cross-validation from CAT. However, this result does not imply that CAT is overparameterized, as suggested by (*19*), without performing tests to validate their assertion. Certainly, CAT uses more frequency categories than nCAT60, but the advantage of Bayesian methods is that they allow safely implementing (*53*) more realistic parameter rich models (*23*), that can be expected to fit to the data better. Indeed, goodness- of-fit tests show that, irrespective of its status as a relative best fit model, nCAT60 fits Whelan- strict (PPA-Div_nCAT60_= 18.9) very poorly, with CAT doing much better (PPA-Div_CAT_= 4.13), a predictable result.

It is our opinion that the solution to the root of the animal tree is not going to be found through a comparison of the number of models (or analyses) supporting Porifera- or Ctenophora-sister. The problem we must solve is why the relative best fit model for the relevant datasets (be that CAT, CAT-GTR, or nCAT60), switch support between Porifera- and Ctenophora-sister as factors such as outgroup sampling or coding strategy are changed. Our hypothesis is that instability under the relative best fit model follows from the fact that also these across-site compositional heterogeneous models struggle to adequately describe these datasets in their amino acid form (*5*). In such situations recodings can improve fit and accuracy, and recoded analyses performed under CAT, CAT-GTR and nCAT60 (see above), in most cases, reject Ctenophora-sister. We can only conclude that, at the least at time of writing, Ctenophora-sister is only clearly supported by analyses of amino acid datasets performed using models that fail to fit the data, a dubious line of evidence.

## Conclusions

Recodings have been widely applied to investigate some of the trickiest nodes in the tree of life, from the root of the animal (*5–7*) to that of plants (*8*), Bacteria (*10*), and Archaea (*9*), with more than 100 papers recently published that used these methods as a line of evidence (*41*). Here we show, using simulations, that recodings are indeed an effective tool to improve phylogenetic accuracy when models do not fit the amino acid data. While a model that does not have a good fit to the data may still infer a correct tree, our simulations show that accuracy significantly improves when models that fit the data well are used. Real datasets assembled to resolve tricky nodes in the tree of life usually display high levels of compositional heterogeneity, and generally fail to be adequately described using available compositionally-heterogeneous models (*5*). While the development of models that could adequately fit arbitrary levels of across-site and across-lineage compositional heterogeneity would be welcome. It is clear that such models are not yet on the horizon – but see (*50*) for some interesting recent progress. Until such models are developed the analysis of recoded data will remain a useful line of evidence in the analyses of the heterogeneous datasets generally associated with tricky nodes in the tree of life.

## Material and Methods

We tested the efficacy of exchangeability-informed recodings using simulations and real data, performing a variety of tests that are summarized in Fig. S2.

### The simulated data

We used the simulated dataset of (*18*), where the target tree used to generate the data assumes either Ctenophora-sister or Porifera-sister to be true (Fig. S1). For each target topology, the 100 (30,000 sites and 97 taxa) datasets of (*18*) were evolved under CAT-LG+G, an across-site rate and compositional heterogeneous model. The original alignments were subsampled to generate datasets with different numbers of characters and taxa. We generated datasets of 1,000, 5,000, 10,000 and 30,000 sites and 20 taxa; as well as datasets of 30,000 sites with 4 and 20 taxa. The number of taxa and characters used in our datasets was chosen to achieve multiple goals. First, we wanted to test the use of recodings using genomic-scale datasets (our 30,000 sites) comparable to those used in real studies. In addition, we wanted to maximize comparability with (*41*) that used 20 taxa and 1,000 to 5,000 sites alignments. Furthermore, we wanted to test the effect of alignment size (1,000 to 30,000 sites), and taxon sampling density (4 to 20-taxon datasets). When subsampling characters from the datasets of (*18*), we made sure that each character was sampled only once in each dataset, and we retained taxa in such a way as to continue to include all key lineages necessary to define the two target topologies (see Fig. S1). Taxon subsampling was fixed so that all our 20, 10 and 4 taxon datasets include the same terminals (Fig. S1). A comparison between our results and those of (*18*) indicated that, by subsampling taxa, we made the inference problem harder (*58*), as success rates for our 30,000 amino acid datasets is lower than that of (*18*) under the same models. This is an advantage for our study as it allows comparing the success rate of amino acids and Dayhoff-6, in more challenging conditions. The datasets of (*18*) were generated using CAT-LG+G. Accordingly, they are only affected by across-site compositional heterogeneity. This is not a problem because exchangeability- informed recordings are expected to reduce both across-site and across-lineage compositional heterogeneity, see (*5*).

#### Comparing simulated and real data

We first investigated whether the across-site compositional heterogeneity in our datasets was comparable to that of real data. This is necessary to understand the extent to which the analysis of the simulated data can inform real world studies. To achieve this goal, we generated 100 datasets of 1,000 characters and 20 taxa each, subsampling the dataset of (*49*). These 1,000 sites datasets were then compared against the 20 taxa and 1,000 sites datasets generated subsampling the simulated datasets of (*18*) – under Porifera-sister only. PPA of site-specific amino acid diversity (PPA-Div (*44*)) was used to test whether the across-site compositional heterogeneity of these datasets could be adequately modeled using GTR+G (referred to as GTR in the main text), a compositionally homogeneous model. Z-scores, the number of standard deviations separating the heterogeneity of the original dataset from the average heterogeneity of a set 100 posterior predictive datasets simulated under the tested model (in this case GTR) were used to measure the ability of GTR to adequately describe the data (*5*). As GTR is compositionally homogeneous, and cannot accommodate across-site compositional heterogeneity, if the dataset generated subsampling the alignments of (*18*) are across-site compositionally heterogeneous, we expect that GTR will fail to adequately model them. Furthermore, If the heterogeneity of the simulated data is comparable to that of real world data, Z-scores from these datasets should be comparable to those obtained from datasets subsampled from (*49*).

Z-scores within the interval included between 2 and -2 indicate that the hypothesis that the model fits the data cannot be rejected (*5*). Z-scores outside this interval could be taken to indicate that the hypothesis that the model fits the data is rejected. However, these boundaries are arbitrary, as the fit of a model achieving a Z-score of 1.99 and that of a model achieving a Z-score of 2.1 is essentially the same. We prefer a more relaxed interpretation of Z-scores and instead of discussing models as being rejected or not, we say that when -2< Z <2 the model fits the data well, or that it adequately describes the data. When 2< |Z| <5 we consider the fit to vary from fairly good (values close to 2) to fairly poor (values close to 5). As a rule, it seems safe to assume that when 5< |Z| < 10 the fit can be considered poor, with |Z|> 10 indicating a very poor fit of the model to the data. We acknowledge that our boundaries are arbitrary. However, they are in our opinion conservative.

All PPA analyses performed to compare real and simulated data were performed in Phylobayes MPI (*59*) v1.8 (One run, total number of cycles= 1,000; burnin= 500; subsampling frequency= 5). .

#### Analyses of the simulated datasets

For all our analyses we recoded the data using Dayhoff-6 (*60*), which was chosen because it is the most widely used exchangeability-informed recoding (see Fig. 1 for details).

30,000 sites and 20 taxa datasets (assuming both Porifera and Ctenophora-sister to be true) were analyzed under a variety of models GTR+G, nCAT10-F81+G, nCAT30-F81+G, nCAT60-F81+G, nCAT120-F81+G, nCAT240-F81+G, CAT-F81 and CAT-GTR+G (also referred to as GTR and nCAT10 to CAT-GTR in the main text). 1,000 to 10,000 sites datasets were analyzed under nCAT10 only, to test whether the performance of recodings is alignment- size dependent. 30,000 sites and 20 taxa datasets were analyzed using all the above-mentioned models. Datasets of 4 and 10 taxa and 30,000 sites were analyzed using nCAT10 only to test whether the performance of recodings depends on the density of taxon sampling. The model used for the analyses using 1,000 to 10,000 site alignments and 4 and 10 taxon alignments (nCAT10) was selected *a posteriori* (see main text), as it emerged from the analyses of 30,000 sites and 20 taxa, that nCAT10 maximizes our ability to discriminate between results of Dayhoff-6 and amino acid analyses with our simulated alignments.

All Bayesian analyses were performed in Phylobayes MPI (*59*) v.1.8 (total number of cycles= 2,000; burnin= 1,000; subsampling frequency= 10). For each dataset a single run was completed to limit the computational burden and minimize pollution, as customary, see (*15, 19*). However, to make sure that 2,000 cycles were enough, for the nCAT10 analyses, we performed two more independent runs using 10,000 cycles, testing congruence after 2,000 and 10,000 cycles.

An analysis was deemed successful if it recovered the target tree with a PP≥ 0.5. To make sure that using a PP= 0.5 cutoff to define success did not bias our results, under nCAT10, we also tested the effect of defining a success using a much more stringent PP=0.95 threshold. In addition, ML analyses were performed in RAXML (*61*) under GTR for the 30,000 sites and 20 taxa datasets, with 200 bootstrap replicates, to make sure that using Bayesian analysis did not bias our conclusions.

For the 30,000 sites datasets, PPA-Div were completed, for each considered model (total number of cycles= 2,000; burnin= 1,000; subsampling frequency= 10). These tests were performed for both the amino acid and the Dayhoff-6 recoded data in Phylobayes (under Porifera-sister only). We performed some tests (not shown) to make sure that PPA-Div results did not change using chains from the analyses performed with datasets assuming Ctenophora-sister to be true and found that the results did not change. Z-scores (see above) were used to report the results of all PPAs.

#### Amino acid jackknife experiment

To test whether the results of analyses performed using Dayhoff-6 might be driven by an information loss induced when the data are recoded, we performed an experiment where we assumed that every substitution in a dataset is phylogenetically informative. This is an overestimate of the true phylogenetic signal of a dataset (see main text). The experiment represents a worst-case scenario because in a situation of this type, every substitution masked by Dayhoff-6 will reduce phylogenetic signal. If all substitutions in a dataset were phylogenetically informative the phylogenetic signal of a dataset could be estimated as the tree length for the considered dataset. We thus estimated the tree length of our amino acid and Dayhoff-6 datasets and calculated the percent of substitutions observed in the amino acid dataset that are masked by Dayhoff-6 – this turned out to be 49.23%. We randomly stripped 49.23% of the sites in our amino acid datasets (to simulate a loss of information comparable to that that Dayhoff-6 would induce in our worst- case scenario). The tree length of the 49.23% jackknifed datasets was re-estimated to quantify whether the number of substitutions in these datasets was now comparable to that in Dayhoff- 6 datasets. The results showed that the jackknifed datasets and Dayhoff-6 datasets had a comparable number of substitutions. We then proceeded to analyze the jackknifed datasets under nCAT10, estimating PPA-Div values, TSR and comparing these values against those obtained under Dayhoff-6, and for the full-length amino acid datasets. All parsimony analyses were performed using PAUPv4.168 (*62*) – Parsimony trees were inferred from 1,000 heuristic searches with the multree option turned off. Tree lengths were then estimated using the Pscore command. Parsimony analyses used the 30,000 sites and 20 taxa datasets.

#### Randomly generated recodings experiment

A perl script (Recodings_generator.pl available: https://bitbucket.org/bzxdp/giacomelli_et_al_2022_recodings/src/master/) was written to randomly generate recodings based on the Dayhoff-6 amino acid partitioning scheme: one 5-state bin, two 4-state bins, two 3-state bins and one 1-state bin. This script was used to randomly generate 1,000 recodings. The similarity of these recodings to Dayhoff -6 was then estimated using a second script (Recodings_comparator.pl available: https://bitbucket.org/bzxdp/giacomelli_et_al_2022_recodings/src/master/). The latter uses intersections of sets to estimate the percent similarity between bins in randomly generated recodings and their exchangeability-informed parent recoding. The same script was used to compare the four randomly generated recodings of (*19*) to SR6 (*31*), their parent exchangeability-informed recoding.

We randomly generated 100 recodings of 0% similarity to Dayhoff-6 (0%-recodings), and 100 recodings of 90% similarity to Dayhoff-6 (90%-recodings). To do this, we first generated 20,000,000 recodings using Recodings_generator.pl. After that, we used Recodings_comparator.pl to identify 100 recodings of specified similarity to Dayhoff-6.

Each (30,000 sites and 20 taxa) dataset was randomly associated to one of the randomly generated 0%- and 90%-recodings. We estimated the tree length for the randomly recoded (0% and 90%) datasets using PAUPv4.168 (same settings as above). The tree lengths of these randomly recoded datasets were compared against those of amino acid and Dayhoff-6 recoded datasets to test whether Dayhoff-6 silences more substitutions than randomly generated recodings (see main text). After that PPA-Div scores and TSRs were calculated (under nCAT10) for the 0%- and 90%-recoded datasets and compared against those obtained using amino acids and Dayhoff-6 (see main text).

#### Analysis of real data (Whelan-Strict)

We applied our approaches to Whelan-strict (*17*), the dataset constituting the main focus of (*19*). This dataset was analyzed as an amino acid alignment and as a 0%-recoded, 90%-recoded and Dayhoff-6 recoded dataset, under the model preferred by (*19*), nCTA60. PPA-Div and PPA-Max values were estimated for each recoding. PPA-Max (*14*) evaluates whether a model can describe the maximal compositional heterogeneity observed across the taxa of a dataset. This test was not used for our simulated data because the model used to simulate the datasets was not across-lineage compositionally heterogeneous, and thus the simulated alignments were not across-lineage compositionally heterogeneous. However, real datasets are expected to be both across-site and across-lineage compositionally heterogeneous, and hence the ability of the considered model to describe both forms of heterogeneity need to be tested for Whelan-Strict. Support values obtained for both Porifera-sister and Ctenophora-sister under all recoding strategies considered, were plotted and compared. In addition, we compared these values against the support values (not the SRs) obtained (in our simulations), when Ctenophora-sister and Porifera-sister were true. We plotted the values obtained under nCAT10, because this model achieved PPA-Div scores comparable to that achieved by nCAT60 for Whelan-strict, suggesting a comparable fit of nCAT10 with our simulated data and nCAT60 with Whelan- Strict.

## Acknowledgments

We would like to thank Herve Philippe and Gert Wörheide for suggestions on previous iterations of this work. We would also like to thank all the members of the Pisani, Donoghue and Williams labs, at the University of Bristol, for discussing these ideas.

## Funding

European Union’s Horizon 2020 research and innovation program under the Marie Skłodowska-Curie grant agreement (764840) IGNITE (M.G./D.P).

NERC GW4+ Doctoral Training Partnership studentship from the Natural Environment Research Council (S100413) (M.E.R/D.P.).

Beatriu de Pinós (Generalitat de Catalunya, 2017-BP-00266) and Juan de la Cierva Incorporación fellowships (Ministerio de Ciencia e Innovación, IJC2018-035237-I) (J.L.F).

Royal Society University Research Fellowship (UF160226) (R.F.)

## Author contributions

Conceptualization: D.P., M.G. Methodology: M.G.,D.P.

Data curation: M.G.,D.P. Writing-original draft: M.G. D.P.

Writing-review & editing: M.G.,D.P.,M.E.R.,J.L.F.R.F. Supervision:D.P.

## Data availability

All the data, material and custom scripts used in this study available in the Bitbucket repository: https://bitbucket.org/bzxdp/giacomelli_et_al_2022_recodings/src/master/

